# Multiple domains of the integral KREPA3 protein are critical for the structure and precise functions of RNA Editing Catalytic Complexes in *Trypanosoma brucei*

**DOI:** 10.1101/2023.04.19.537538

**Authors:** Brittney Davidge, Suzanne M McDermott, Jason Carnes, Isaac Lewis, Maxwell Tracy, Kenneth D. Stuart

## Abstract

The gRNA directed U-insertion and deletion editing of mitochondrial mRNAs that is essential in different life cycle stages for the protozoan parasite *Trypanosoma brucei* is performed by three similar multi-protein catalytic complexes (CCs) that contain the requisite enzymes. These CCs also contain a common set of eight proteins that have no apparent direct catalytic function, including six that have an OB-fold domain. We show here that one of these OB-fold proteins, KREPA3 (A3), has structural homology to other editing proteins, is essential for editing and is multifunctional. We investigated A3 function by analyzing the effects of single amino acid loss of function mutations most of which were identified by screening bloodstream form (BF) parasites for loss of growth following random mutagenesis. Mutations in the ZFs, an intrinsically disordered region (IDR) and several within or near the C-terminal OB-fold domain variably impacted CC structural integrity and editing. Some mutations resulted in almost complete loss of CCs and its proteins and editing whereas others retained CCs but had aberrant editing. All but a mutation which is near the OB-fold affected growth and editing in BF but not procyclic form (PF) parasites. These data indicate that multiple positions within A3 have essential functions that contribute to the structural integrity of CCs, the precision of editing and the developmental differences in editing between BF and PF stages.

## INTRODUCTION

Mitochondrial mRNAs in *Trypanosoma brucei,* the causal agent of Human African Trypanosomiasis (aka sleeping sickness), the related *T. cruzi* and *Leishmania spp*. pathogens and other kinetoplastids undergo post-transcriptional maturation by RNA editing (Read et al. 2016). The editing generates mature functional mRNAs from transcripts by the insertion and deletion of U-nucleotides as specified by small guide RNAs (gRNAs) (Koslowsky et al. 1990; Pollard et al. 1990; Riley et al. 1994). Some mRNAs undergo quite limited editing, e.g. the insertion of 4 or 34 Us in COII and CYb mRNAs, respectively, whereas editing essentially recodes the sequences of other mRNAs by extensive U insertion/deletion e.g. of +447/-28 and +547/-41 Us respectively in mRNAs for ATPase 6 and COIII oxidation-phosphoryation complex proteins and +132/-28 in RPS12 mitoribosomal protein mRNA (Benne et al. 1986; Feagin et al. 1987; Feagin et al. 1988; Bhat et al. 1990; Read et al. 1992). In addition, the editing of several transcripts differs between the life cycle stages of *T. brucei* in parallel with its metabolic and developmental differences. Edited cytochrome subunit mRNAs are abundant in the insect midgut stage procyclic form (PF) parasites which generate energy via oxidative phosphorylation whereas these edited mRNAs are dramatically reduced in the mammalian bloodstream form (BF) parasites that produce energy via glycolysis (Panigrahi et al. 2008). Thus, this differential editing adapts this pathogen to the disparate environments of the midgut of its tsetse fly vector vs. the mammalian bloodstream (Feagin et al. 1987). The mechanisms responsible for this developmental difference are unknown.

RNA editing is catalyzed by three similar multiprotein Catalytic Complexes (CCs) that have different editing site specificities. Each round of editing is initiated by cleavage of an editing site (ES) by an endonuclease. Following cleavage, Us are either inserted by a TUTase or removed by an ExoUase. Lastly, the ES is re-ligated by an RNA ligase. Each CC contains a common set of twelve proteins including TUTase, ExoUase and RNA ligase enzymes, as well as eight proteins that have no apparent catalytic function. Mutually exclusive pairs of endonuclease and partner proteins functionally delineate each of the three CCs, one of which also contains an additional ExoUase (Schnaufer et al. 2003; Worthey et al. 2003; Carnes et al. 2017). KREPA3 (A3) is one of the non-catalytic proteins that is common to all three CCs. It contains two C2H2 zinc fingers motifs (ZFs), a C-terminal OB-fold motif, several intrinsically disordered regions (IDRs) but no other motifs that are related to known protein domains or predicted structures (Panigrahi et al. 2001; Schnaufer et al. 2010).

A3 is essential for cell growth and functions in RNA editing in both BF and PF life cycle stages as has been shown by the loss of parasite viability and RNA editing upon A3 expression knockdown by RNAi or in conditional null (CN) cell lines (Guo et al. 2008; Law et al. 2008; Guo et al. 2010; McDermott et al. 2015b). Loss of A3 expression also affects CC structural integrity, albeit to a greater extent in BFs than in PFs as shown by the retention of CCs in PF but not BF A3 CN cells (Brecht et al. 2005; Guo et al. 2008; McDermott et al. 2015b). Mutation analyses showed that the N-terminal ZF (NTZF) and the more C-terminal ZF (CTZF) domains are both required for parasite viability and RNA editing in BFs whereas the latter ZF is not for required in PF (Guo et al. 2008; Guo et al. 2010; McDermott et al. 2015b). Thus, the two A3 ZFs impact CC structure and function somewhat differently between these two life-cycle stages.

The C-terminal region of A3 contains a prominent OB-fold domain with a β-barrel comprised of five strands (β_1_-β_5_). Like other OB-fold proteins, β_1_and β_2_, β_2_ and β_3_, and β_4_ and β_5_ are connected by loops L_12_, L_23_, and L_45_, respectively, while strands β_3_ and β_4_ are connected by an α-helix, C_34_ (Theobald et al. 2003; Horvath 2011). The region adjacent to the N-terminal strand of the OB-fold is distinct from the corresponding region of the OB-folds of the other five CC proteins (A1,2, 4-6) (Park and Hol 2012). Previous studies have described the structure of the A3 OB-fold and its proximity to other proteins in the CC complex (Schnaufer et al. 2010; Park and Hol 2012; McDermott et al. 2016). Yeast two-hybrid and cross-linking-MS studies have shown that the A3 OB-fold interacts with B5, another non-catalytic CC protein and is proximal to several other proteins in CCs including the N1 deletion endonuclease (Schnaufer et al. 2010; McDermott et al. 2016), N2 insertion endonuclease and the five other OB-fold containing proteins (Schnaufer et al. 2010; McDermott et al. 2016). Previous studies have shown that the OB-fold can interact with RNA in vitro and has RNA chaperone activity (Brecht et al. 2005; Voigt et al. 2018). Structure determination of recombinant A3 and A6 co-crystals identified interactions between A3 and A6 OB-folds within the crystal lattice (Park and Hol 2012; Park et al. 2012). Thus, the A3 OB-fold likely plays a structural role in CCs via its interactions with other CC proteins including close interactions with A1-A6 as shown by cross-linking studies (Schnaufer et al. 2010; McDermott et al. 2016; Voigt et al. 2018). However, the specific functional roles of A3 *in vivo* in CC structure and editing and how they may differ between BF and PF stages are unknown (Schnaufer et al. 2002; McDermott et al. 2015b).

We report here the identification of multiple single amino acid mutations that result in loss of function (LOF) in BFs and in one case in PFs. Most of these substitutions mapped to the two ZFs, an IDR in a region with no predicted structure or in or adjacent to the OB-fold. Exclusive expression of some of these mutant alleles resulted in loss of all three CCs and editing in BFs but not in PFs except in one case where CCs and editing were reduced but not eliminated in PFs. Other mutations had lesser effects on CC structure and altered but did not eliminate RNA editing and some of these mutations had differential effects on CCs and editing that were generally more impactful in BFs than in PFs. These results indicate that A3 interacts with multiple proteins in the three CCs and contributes to their structural organization and integrity. The results suggest that various A3 domains identified by these mutations have specific functions among the numerous events that occur during the complicated process of RNA editing. These functions likely occur via specific molecular interactions that may differ among the three CCs which possess different editing site specificities. The differential effects of the mutations between BF and PF cells imply that some of these A3 domains function differently between these stages despite the indistinguishable protein compositions of CCs.

## RESULTS

### Mutations that affect growth of BFs are in multiple protein domains

To elucidate the role of A3 in editing we tested 634 BF CN cell lines that had been cloned in wells by dilution after transfection with a library that contained ∼10,000 unique full length randomly mutated A3 alleles and identified 136 wells with growth defects by replica plating in the presence or absence of tet as previously described (Table S1) (McDermott et al. 2015a). PCR and sequencing of A3 from 59 wells that had strong growth defects identified 14 mutations of interest from sequences that 1) had a single amino acid substitution, the same substitution in multiple clones that had multiple mutations, or a substitution that likely would have a functional consequence based on structural analyses, and 2) which reproduced the growth defect upon exclusive expression in cells in which the mutation was independently reconstructed by site-directed mutagenesis (Fig. 1A). Three single substitutions (Q299H, L262P and T315A) did not reproduce the growth defects upon reconstruction and thus these residues are not critical for A3 function in BFs and may have originated from wells in which the screened cells were not clonal. This number of confirmed single amino acid substitutions that affect A3 function is similar to that identified in B5 (9), B6 (12), B7 (10), and B8 (9) by similar methods (McDermott et al. 2015a; Carnes et al. 2022). We also constructed cell lines with double amino acid substitutions in the more N and C-terminal ZFs (NTZF and CTZF) with an N-terminal V5 tag for comparison with previous constructs carrying the same mutations (Guo et al. 2008; Guo et al. 2010) (Table S2).

**Figure 1:**
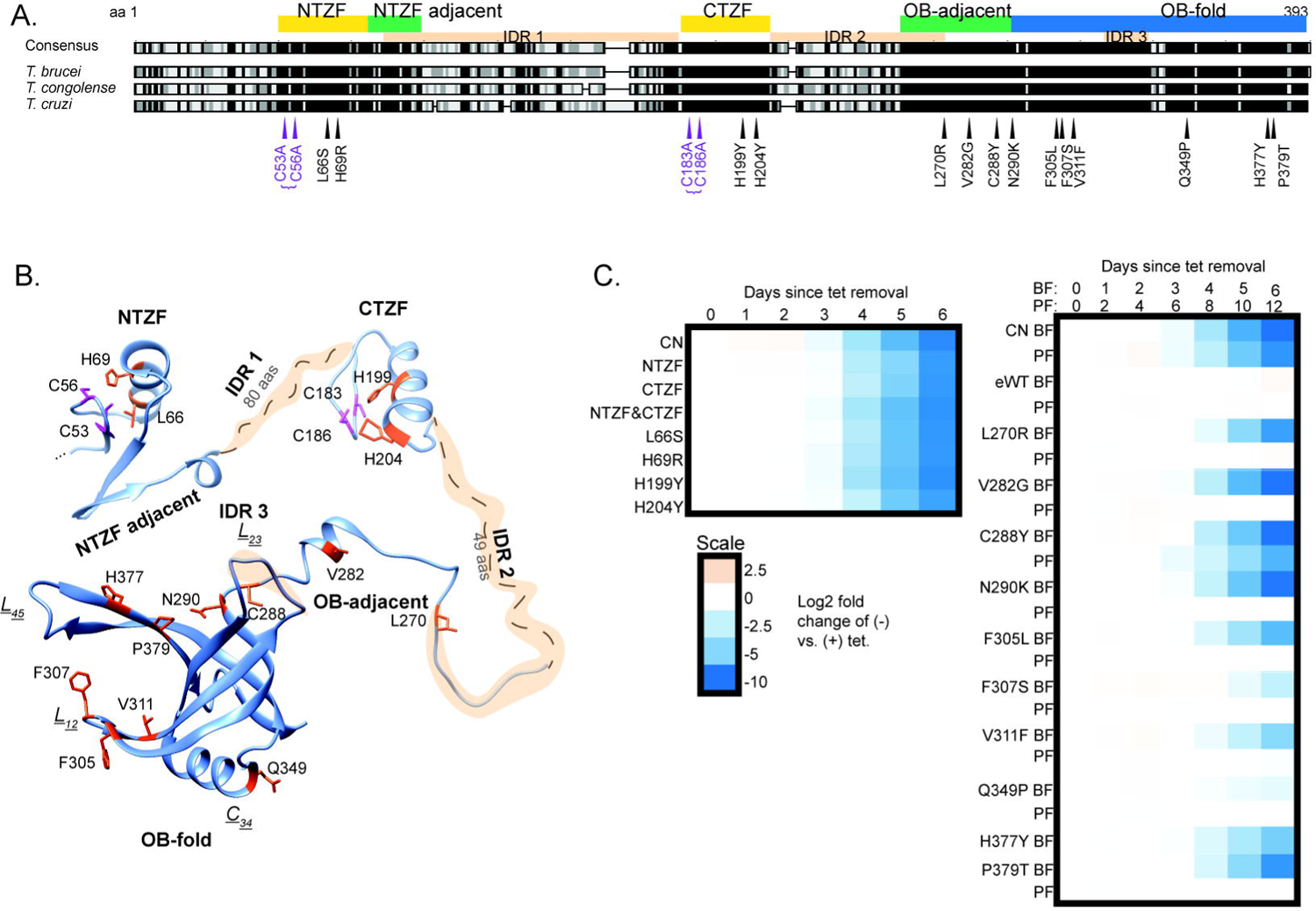
Loss of function mutations in A3 and their effects on growth. **A**: Locations of single and double (bracketed) amino acid substitutions that resulted in loss of growth in BF *T. brucei* when exclusively expressed. Substitutions are mapped against a sequence alignment of three Trypanosoma species; black indicates amino acid similarity. The locations of the NTZF and CTZF (yellow), The OB-fold (blue) and the regions adjacent to these (green) are indicated as are three IDRs (tan). **B**: Location of the substitutions in a composite A3 structure of a high confidence AlphaFold structural prediction (veb8-A) and crystal structure of the OB-fold (PDBID: 4DNI, Park and Hol 2012). Single amino acid substitutions and double C➔A substitutions are in magenta. **C**: Effect on growth after A3 repression (CN) or of exclusive expression of A3 (i.e. -tet) with substitutions in or near the ZFs (left) or in or near the OB-fold (right) over 6 days in BFs and 12 days PFs (see methods for details).

The confirmed LOF mutations localized to the two ZF domains and to various positions near or within the OB-fold (Fig. 1A). They were mapped to the partial A3 structure that had been determined by x-ray crystallography and a full-length A3 structure that was predicted using AlphaFold (Park and Hol 2012; Jumper et al. 2021) (Fig 1B). The double mutations in the ZFs map to regions that are adjacent to a helix that is in each ZF, the single substitutions map to similar positions within these small helices in each ZF, which except for L66S are likely to impact coordination of one or both Zn^2+^ ions coordinated by C2H2 ZF domains similar to the two double mutations. The region adjacent to the NTZF has predicted β-strands that contains many charged lysine and glutamate residues. The features of these ZF domains imply that they participate in RNA and/or protein interactions. The other single mutations mapped to a region adjacent to the N-terminal side of the OB-fold which includes one of the predicted intrinsically disordered (IDR) inter-domains and a small helical region and to multiple positions within the highly structured OB fold (Fig. 1B). The single and double mutations in the ZFs had substantial impacts on growth of the BFs (Fig 1 C); the single ZF mutations were not tested in PFs but double mutations in the NTZF but not the CTZF substantially inhibit growth in PFs (McDermott et al. 2015b). The mutations that mapped to the region that is adjacent to or in the first strand of the OB-fold (V282G, C288Y and N290K) or the nearby disordered region (L270R) that was previously predicted (Park and Hol 2012) also had substantial impacts on growth (Fig. 1C). The mutations that mapped to the core of the OB-fold β-barrel structure or in or near loops L_12_ and L_45_ (F305L, F307S, V311F, Q349P, H377Y and P379T) had lesser effects on growth. All mutations throughout the OB-fold and the nearby region except C288Y and C53A/C56A mutations in the more N-terminal ZF resulted in growth defects in BFs but not in PFs indicating that these domains may function in the differential editing that occurs between these life cycle stages (Fig. 1C) (McDermott et al. 2015b).

### Mutations of A3 affect CC abundance and integrity

Westerns of Blue Native (BN) and SDS PAGE analysis of lysates from the reconstructed A3 mutant BF and PF cell lines with LOF mutations of A3 revealed various effects on CC structural integrity and abundance of CCs and their proteins (Fig. 2). The ∼1MDa complexes, four CC proteins and tagged A3 protein were evident in cells that exclusively express WT A3 (eWT) from the tubulin locus using mAbs for A1, A2, A3 and L1 CC proteins, the V5 tag on A3 and mt HSP70 that was used as a loading control. The parental CN cells, which do not have the tagged A3 allele, lacked CCs and the four CC proteins when A3 expression from the rDNA locus was repressed (-tet). This shows both the stringent loss of A3 protein during exclusive expression, i.e. -tet, and that A3 is required for the presence of CCs and these CC proteins. Notably, A3 consistently runs with a slightly smaller apparent size in the L270R mutant in both BF and PF cells. Genomic DNA sequencing confirmed that the L270R allele is full length, indicating that the mutation itself alters its apparent size perhaps due to an added charge from Arg. Conversely, A3 has a slightly larger apparent size in CTZF single H199Y and H204Y and double C183A/C186A mutants in BFs, perhaps due to the coordinated Zn^2+^ ion. We cannot exclude the possibility that these differences may be due to other processes such as post-translational modifications (PTMs). V5-tagged A3 mutants were present in similar cellular amounts except in C288Y in which there was CC loss. Notably, the A3 mutant proteins detected with the V5 mAb were ∼50kDa in size, as expected, whereas the mutants detected with the A3 mAb were ∼42kDa in size. This size difference is consistent with the N-terminal V5-tag being cleaved off upon import into the mitochondrion. A3 detected with the A3 mAb appeared more variable in amount which correlated with the abundance of CCs and the other three CC proteins.

**Figure 2:**
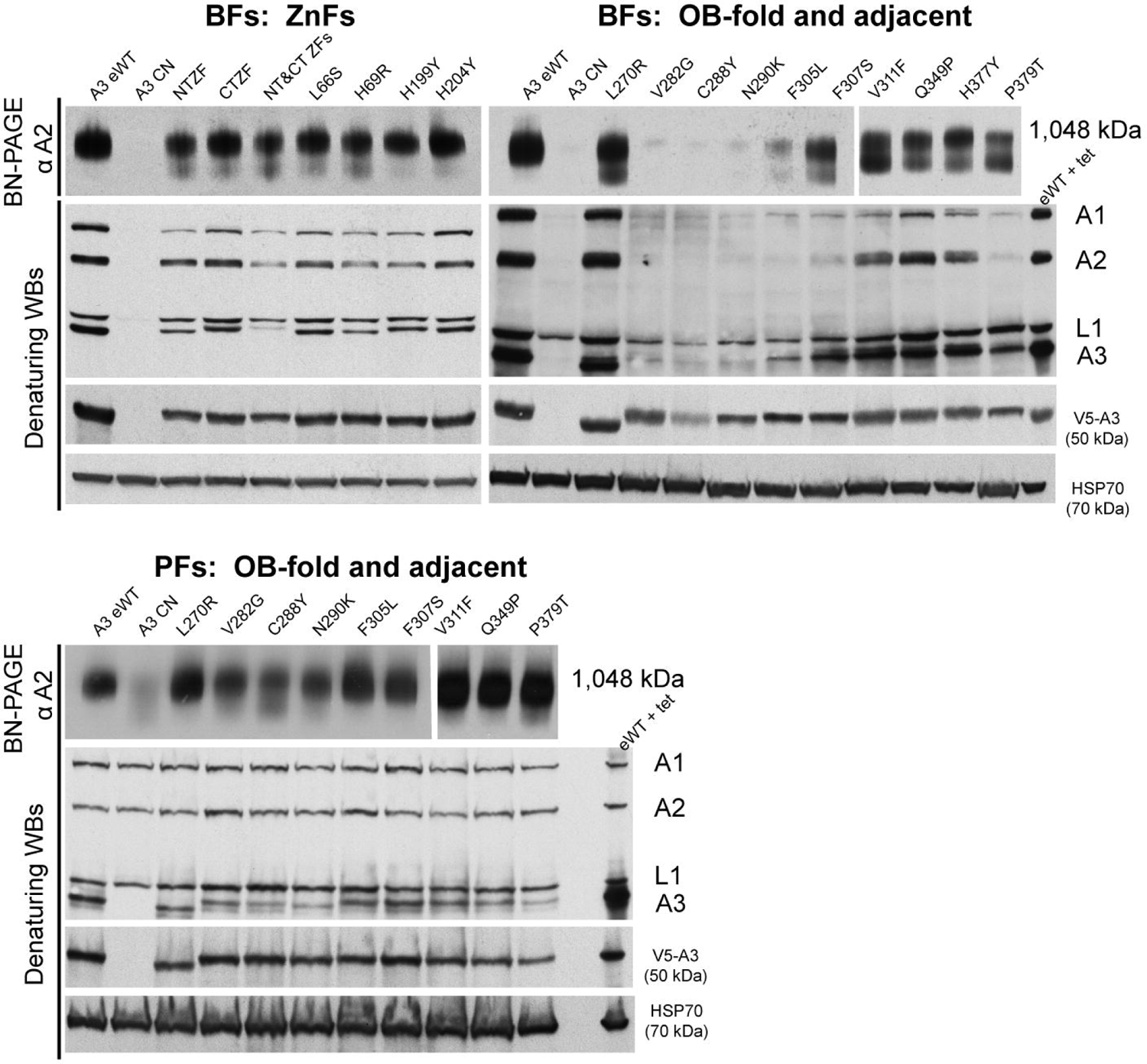
Effects of A3 mutations on Catalytic Complexes. Western analyses of blots analyzing cleared cell lysates after either 48 hrs (BF) or 96 hrs (PF) of exclusive expression of either mutated, wild type, or no A3 (CN) using BN-PAGE (upper panel) and denaturing PAGE (lower panels). The BN-PAGE blots were probed with a mAb for A2 which is present in all CCs, the denaturing PAGE blots were probed with a mAb mixture for four CC proteins or mAbs for the V5-tag on A3, or the HSP70 loading control. Note the greater effect on the CCs in BF (upper) vs. the PF (lower).

The abundance and integrity of CCs and the four CC proteins (A1, A2, A3, and L1) differed between the various mutant cell lines (Fig. 2). The ∼1MDa CCs and the four CC proteins are abundant in all BF cell lines containing mutations in the ZFs but a smear below the ∼1 MDa band suggests some impact on CC structural integrity which is more apparent with mutations in the NTZF with parallel effects on the levels of the four CC proteins assayed. Mutations that are in or near the OB-fold had various effects on the CCs and their proteins (Fig. 2). The V282G, C288Y, and N290K mutants and to a lesser extent the F305L mutant had lost most of the ∼1MDa CCs and the four CC proteins. There were similar but somewhat lower CC and protein losses in the F307S and P379T mutants which contained an ∼800 kDa complex that was more prevalent in V311F and P379T mutants, reminiscent of previously observed subcomplexes (Carnes et al. 2022). There was a general correlation between CC abundance and severity of growth defect for BF cell lines with mutations in or near the OB-fold, as cell lines with fewer intact CCs had greater growth defects (Figs 1 and 2). This was not the case for the ZF or disordered region (L270R) mutants which have abundant ∼1MDa CCs but substantial growth defects (Figs 1 and 2).

Together these data suggest that although the ZFs are important for CC integrity in BFs, they likely play additional functional roles e.g. RNA binding that contribute to the substantial growth defects associated with their mutation. Mutations in or near the OB-fold were also tested in PFs, for which only C288Y has any notable impact on the CCs (Fig. 2). In these cells, a smeared band can be observed on the BN-PAGE gel with sizes ranging from about 800kDa-to 1MDa, indicating CC fragmentation (Fig. 2). V282G and N290K, which have normal growth in PFs, have CCs that might also be fragmented, although the extent of fragmentation is less than in C288Y (Fig. 2). Previous studies have shown that the CTZF is not necessary for growth or CC integrity in PFs, but the NTZF is required for PF growth (McDermott et al. 2015b). Together, these results suggest that different parts of A3, including the OB-fold, disordered region, and at least one ZF, contribute differently to CC structure between life cycle stages.

The effects on CCs which were retained in the BF L66S, L270R, V311F, H377Y, and P379T mutants were examined in more detail by glycerol gradient fractionation. The CCs from eWT, L66S, L270R, and H377Y sedimented with a peak at ∼ 20S (fractions 13-17) as seen in the Western blots of BN and SDS-PAGE blots probed with mAbs for four CC proteins (Figs. 3A and 3B). The blots were also probed with a mAb for RESC1, a protein of the RNA editing substrate complex (SC) with which CCs functionally interact (Carnes et al. 2023), and with a mAb for HSP70 as a loading control (Fig. 3B). The mutations resulted in various reductions in the amounts of CCs per cell as show by the western analyses of lysates of equivalent numbers of cells. Mutations in the NTZF (L66S), the IDR (L270R) cause only minimal, if any, loss of CCs whereas mutations in strands β5 (H377Y and P379T) and β2 (V311F) of the OB-fold had greater CC loss but all retained some ∼20S CCs. The V311F and P379T mutants had visible ∼800kDa bands in fractions 9-11 in the BN-PAGE westerns and preferential loss of the A1 CC protein in the SDS-PAGE blots. In addition, the bulk of the complexes in these mutants are shifted to lower S regions of the gradient where the preferential loss of A1 and A2 is evident in the direct comparisons (Fig. 3C). These results are consistent with the loss of the ∼186kDa A1-containing heterotrimeric insertion subcomplex and retention of a ∼800kDa subcomplex comprised of the L1-containing heterotrimeric deletion subcomplex and other CC proteins (Carnes et al. 2017). The detection of A3 at low S regions of the gradient with anti-V5 antibody but not anti-A3 antibody likely is due to a combination of the effects of the greater sensitivity of the V5 antibody and of the N-terminal V5 tag which may have affected the processing or the MTS (mitochondrial targeting sequence) and transit into the mitochondrion. The mutations had no appreciable effect on the abundance or distribution of the RESC1 component that binds gRNAs (Weng et al. 2008; Hashimi et al. 2009; Aphasizheva et al. 2020) and functionally interacts with CCs (Aphasizheva et al. 2020). Overall these results illustrate the role of A3 in the structural organization of CCs, and the disruption caused by the V311F mutation signifies the importance of a hydrophobic pocket in A3 that includes V311 (Park and Hol 2012) for CC structural integrity.

**Figure 3:**
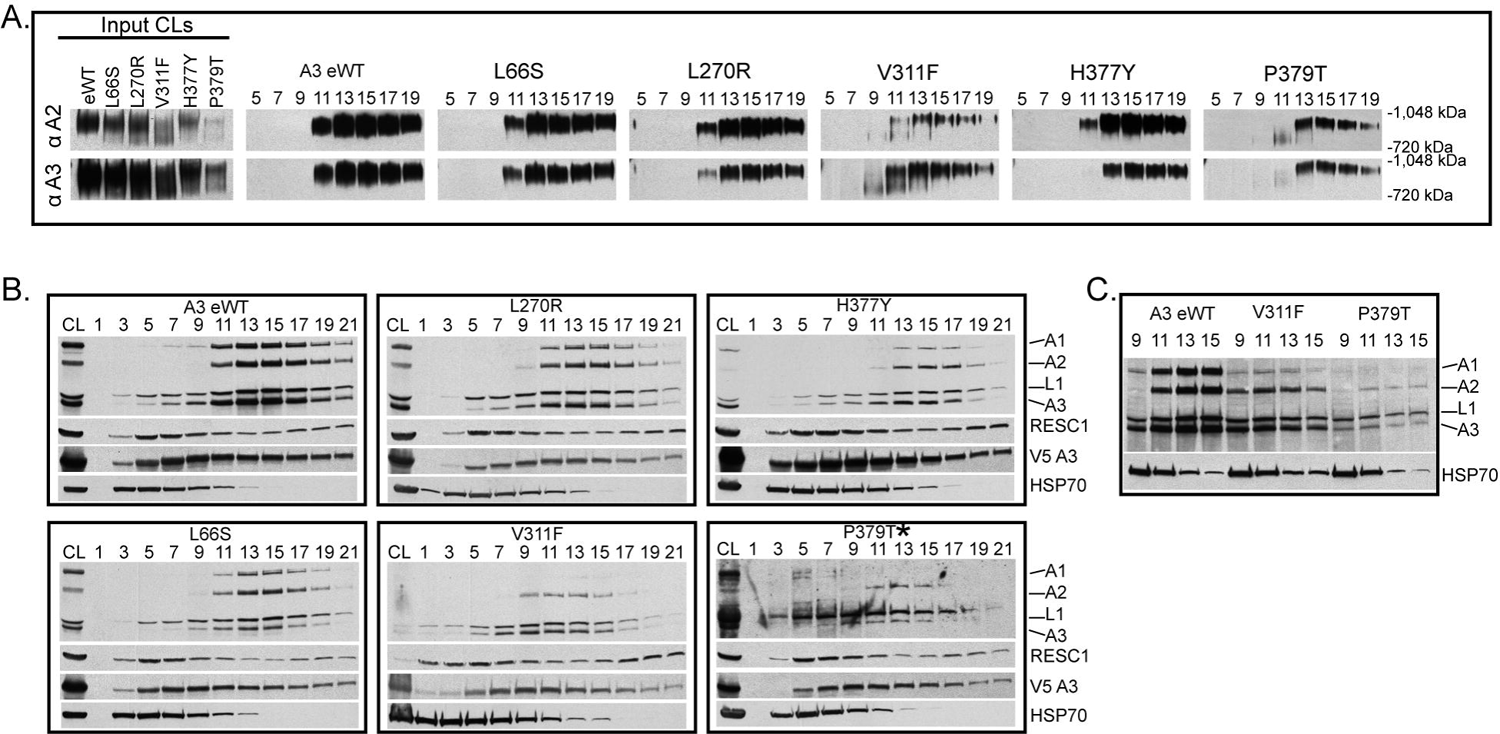
Glycerol gradient analysis of the effects of A3 mutations on Catalytic Complexes. Western analysis of 10-30% glycerol gradient fractions from lysates of the indicated cell lines probed with indicated mAbs as in Fig. 2. **A**: Blots of BN-PAGE of input cleared lysates (CL) and gradient fractions. **B**: Denaturing Westerns of input CL and of glycerol gradient fractions probed as in A. The P379T* blot used supersignal ECL and was exposed longer. **C**: Selected gradient fractions for direct comparisons among eWT, V311F, and P379T.

### Effects of A3 mutations on RNA editing

The effects of selected BF and PF A3 mutations on RNA editing *in vivo* were assessed by high-throughput RT-qPCR using the Fluidigm BioMark system (McDermott et al. 2015b). The abundances of amplicons from pre-edited, edited, and never-edited mitochondrial mRNAs in BF and PF cells that were exclusively expressing WT or mutant A3 alleles were assayed prior to the effect on cell growth relative to those in the corresponding cells in which the tet-regulatable WT A3 allele was expressed (Fig 4A and Table S2). The abundances of most edited transcripts were reduced upon exclusive expression of mutant relative to WT A3 in BFs. The lack of detectable edited CYb mRNA in BFs or 3’ edited ND7 mRNA in PFs reflects their known developmental regulation (Schnaufer et al. 2002; McDermott et al. 2015a). The reductions in edited transcript levels in BF mutant samples generally correlated with the severity of growth defect and CC abundance, e.g., V282G, C288Y, or N290K mutations had the greatest decreases in editing. An exception to this is that the L270R mutation did not impact CC abundance in BFs but had a substantial reduction in edited mRNAs and a moderate growth defect (Fig. 4 compared with Figs. 1 and 2). Strikingly, the effects of these mutations on the abundances of the edited PCR products differed between BF and PF life cycle stages and mirrored the impacts on growth and CC abundance. Only one OB-fold mutation (C288Y) impacted edited mRNA levels in PFs (Fig. 4A). Thus, A3 functions differently in editing between the life cycle stages.

**Figure 4:**
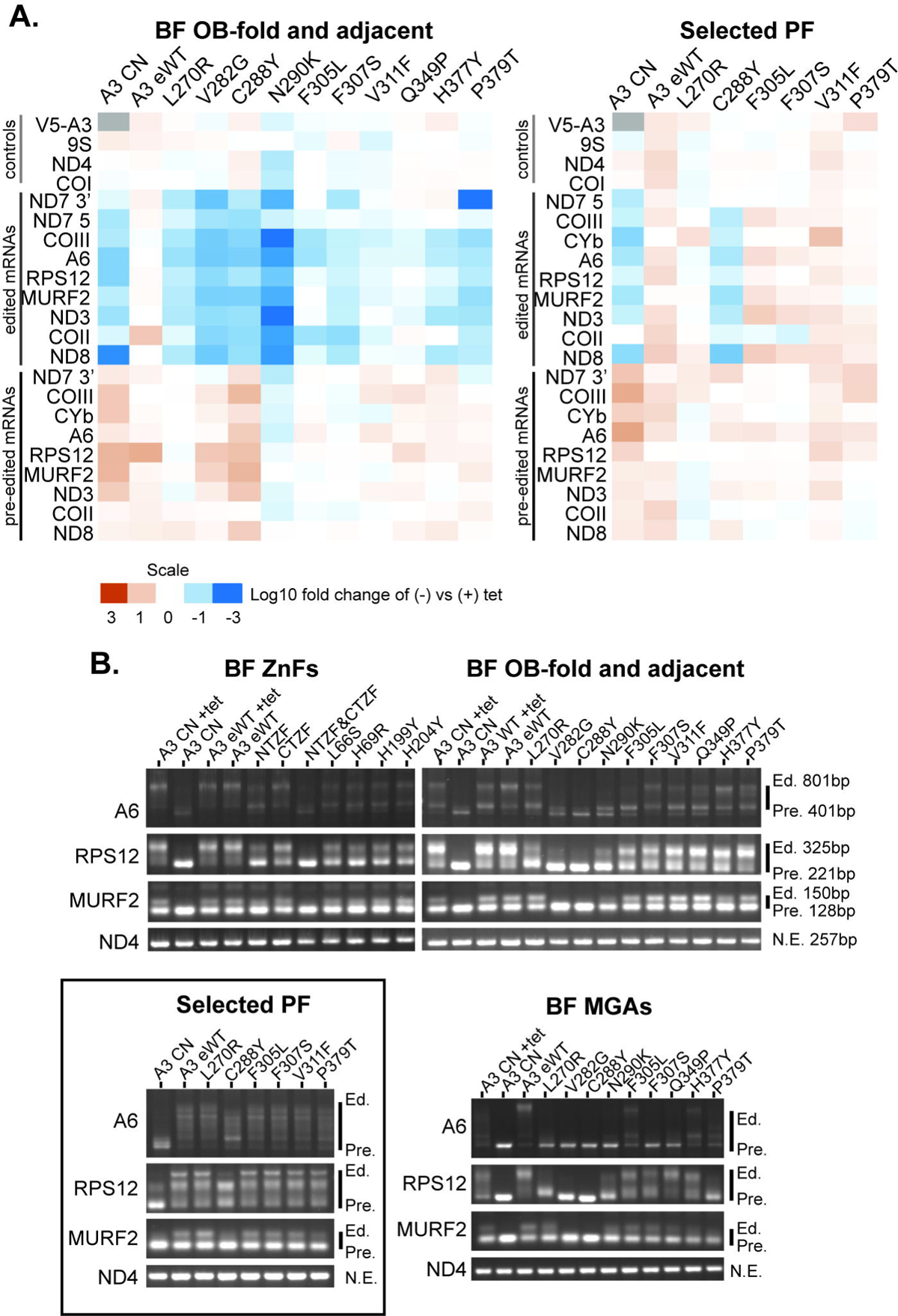
Impacts of A3 mutations on RNA editing. **A**: qPCR measurements of pre-edited and edited mRNAs as indicted in samples from BF and PF cells grown in the presence or absence of tet for 48 and 96 hours, respectively. The levels of RNAs were normalized to the TERT control. Relative abundances of mRNAs (-tet vs + tet) were then determined for each sample, transformed by log(10), and represented by a heat map. **B**: Gel profiles of RT-PCR products with pre-edited (Pre), fully edited (Ed.) sizes as indicted are from cells grown for 48 hours (BF) or 96 hours (BF with an MGA allele or PF) following tet removal. Only minus tet samples are shown unless indicated otherwise. Box: selected PF cell lines that represent different regions of the OB-fold; most mutations do not result in growth defects in PF.

These qPCR analyses provide an overview of targeted pre-edited and edited mRNA amplicon abundances but do not measure partial or anomalous editing resulting from the mutations. We therefore generated profiles of all of these RNAs from BF and PF cells by performing RT-PCR using primers that anneal to the mRNA 5’ and 3’ terminal regions of the mRNAs that do not get edited, resulting in amplification of all products regardless of how much editing occurred. We analyzed ATPase 6 (A6), RPS12, and MURF2 mRNAs which are edited in both life cycle stages and used never-edited ND4 mRNA as a control (Fig. 4B). We resolved the resultant products on an agarose gel to visualize the editing products associated with each exclusive-expression cell line (Fig. 4B). Because edited mRNAs contain more insertion sites than deletion sites, larger products result from more editing, and thus more editing occurred in cell lines with larger products. The A3 CN cell line in both the presence and absence of tet was included as a control. The different mutations resulted in distinct RT-PCR product profiles (Fig. 4B). In BFs, the single L66S, H69R, H199Y and H204Y mutations in the helical regions of either ZF and the double C183A/C186A mutation in the CTZF resulted in A6 profiles that were distinct from the conditional null in the presence or absence of tet, which suggests that some editing occurred but was likely anomalous or incomplete, similar to what has been previously described for double mutation in the ZFs (Guo et al. 2010). A similar result was observed for RPS12 except the single mutants had more pre-edited size product in comparison to the CN +tet. The double C53A/C56A mutation of the NTZF alone or together with the double mutation of the CTZF resulted in greater amounts of pre-edited sized product, which is indicative of reduced editing. These results and those with the double mutation in the NTZF and/or CTZF in BF and PF CN cells that exclusively express a C-terminal TAP-tagged (BF) or untagged (PF) A3 reinforce the role for the A3 ZFs in editing (Guo et al. 2008; Guo et al. 2010; McDermott et al. 2015b).

Mutations in or near the OB-fold had more variable effects on editing which were parallel to their effects of CC structure. The V282G, C288Yand N290K substitutions which resulted in substantial loss of CCs and their proteins (Fig. 2) had predominant RT-PCR products consistent with pre-edited mRNAs or short partially edited products (Fig. 4B). The F305L mutant that had reduced, but not eliminated CC abundance had larger RT-PCR products consistent with partially edited sizes. Similar partially edited products were also observed with F307S, V311F, Q349P, H377Y and P379T mutants that broadly retained CC proteins. Interestingly, the L270R mutant that retained CCs had similar A6 profiles to those in these five mutants but had a prominent RPS12 product that was slightly larger the pre-edited rather than larger products seen in the others (Fig. 4B).

Analysis of the effects of a subset of the OB-fold mutations on editing in PFs by gel analysis of the RT-PCR products of the A6, RPS12 and MURF2 mRNA showed more products that are larger than pre-edited than in BFs which reflect the presence of CCs in PFs. The sizes of the RNAs are similar to those in the eWT with the exception of the C288Y mutants where smaller products are seen (Fig. 4B). Thus, mutations in the OB-fold that were assayed in PFs resulted in different effects on editing than in BFs and indicating functional differences of the A3 OB-fold between these developmental stages.

We transfected a <u>M</u>utant <u>G</u>amma <u>A</u>TP synthase (MGA) allele into the A3 CN cell line and the reproduced some of the mutant cell lines and eWT to enable the survival and continuous growth of BFs in the absence of RNA editing (Figs S1 and 4B) (Dean et al. 2013; Carnes et al. 2017; Carnes et al. 2023). This allowed us to assess possible incomplete development of the effects of the mutations or secondary effects of the mutations, e.g. physiological consequences due to loss of products encoded by edited mRNAs. This resulted in a shift to a greater proportion of products with sizes at or near that of pre-edited mRNA (Fig. 4B). This was especially evident for the V282G and C288Y mutations that lack most CC material (Figs. 2 and S1) and with the longer A6 and RPS12 mRNAs. The other mutations had products that were larger than pre-edited but generally smaller than the largest product in wild type. The L270R mutant retained ample ∼ 1MDa CCs but the RT-PCR product appeared slightly longer than pre-edited (Fig. 2). Overall, these results suggest that editing occurred in mutants that retained CCs albeit insufficient in amount or accuracy to support growth.

To determine the characteristics of the RT-PCR products we sequenced multiple RPS12 PCR products that range in size from ∼221bp (pre-edited) to 325bp (fully edited) that we cloned from cell lines that retained various proportions of ∼1MDa and ∼800 kDa CCs. As shown in the diagram (Fig. 5) these sequences indicated that editing had occurred in these cells and is incomplete compared to fully edited RNA including in WT as is typical. However, in the L270R mutant the extent of editing and the proportion of fully edited insertion ESs were reduced and the proportions of deletion ESs that were not fully edited and of editing in non-edited ESs were increased. In the F307S and P379T mutants the extent of editing was greater than in L270R and there were proportional differences in the various types of ESs from those in WT and L270R but not statistically significant in this survey less than in the WT (Fig. S2). These results show that these mutants perform editing but do so with an accuracy and/or efficiency that is insufficient for cell viability.

**Figure 5:**
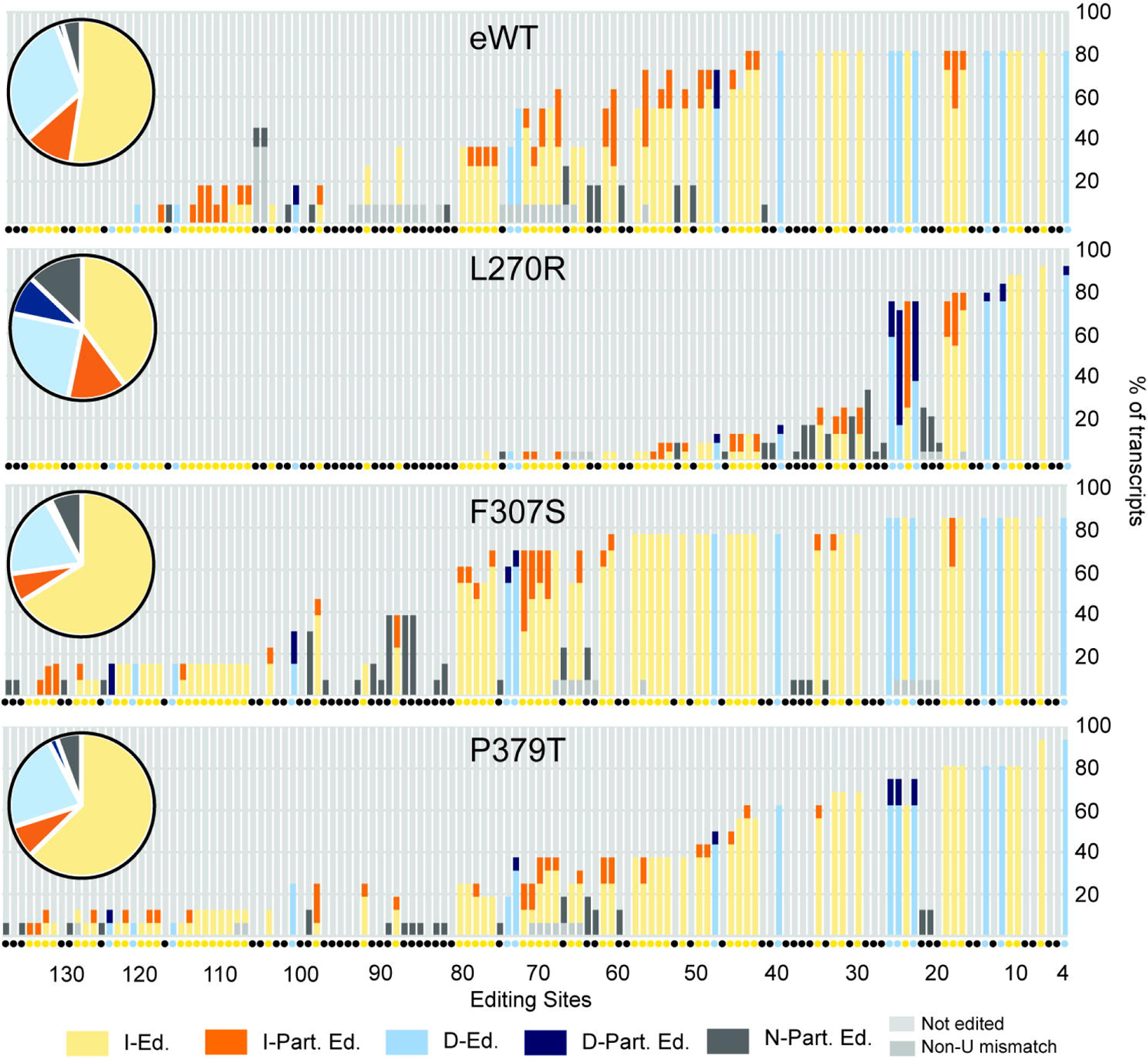
Impacts of A3 mutations in BF cell lines on cloned RPS12 sequences. A summary of the editing of cloned and sequenced RPS12 mRNAs (Fig. S2) is summarized by the bar graph that shows the percent of the type of editing at each ES classified as I-Ed.= fully edited insertion ES (yellow), I-Part. Ed.= partially edited insertion ES (orange), D-Ed.= fully edited deletion ES (light blue), D-Part. Ed.= partially edited deletion ES (dark blue), N-Part. Ed.= partially edited ES that is not edited in fully edited mRNA (dark grey). ESs where no editing was observed (light grey) and ESs impacted by a non-U mismatch (medium grey) are shown. Each type of editing event is shown as a percent of the total for each sample in the pie chart inserts. The colors of the small circles beneath each bar indicate the type of editing at each ES in fully edited mRNA.

### Structural modeling and functional context of the effects of A3 mutations

A Dali search (Holm 2022) for *T. brucei* structural homologs of A3 identified the A1, A2, A4-6 OB-fold containing CC proteins as expected (z-scores of 14.7, 12.6, 10.6, 13.7, 13.4, respectively). It also identified Tb927.9.4810 MPSS5 (z-score 10.0) and Tb927.6.2190 MPSS6 (z-score 11.1) components of the MPsome complex that has TUTase and exonuclease activities and gRNA processing functions and Tb927.10.8220 (z-score 11.1) an uncharacterized protein that has been observed in isolations via T1, DSS1, and MPSS2 pull-downs and whose gene is adjacent to A2 (Fig. 6A) (Penschow et al. 2004; Aphasizheva and Aphasizhev 2016; Suematsu et al. 2016; Aphasizheva et al. 2020). All of these proteins have predicted OB-folds similar to A3, with amino acids cognate to those identified as critical for A3 function (Tables S3 and S4). Additionally, MPSS5 has a C2H2 ZF similar to A3, while MPSS5 and MPSS6 have predicted helical domains and regions with no predicted strictures that may contain IDRs, all features that imply functional interactions with RNA and/or protein (Fig. 6A).

**Figure 6:**
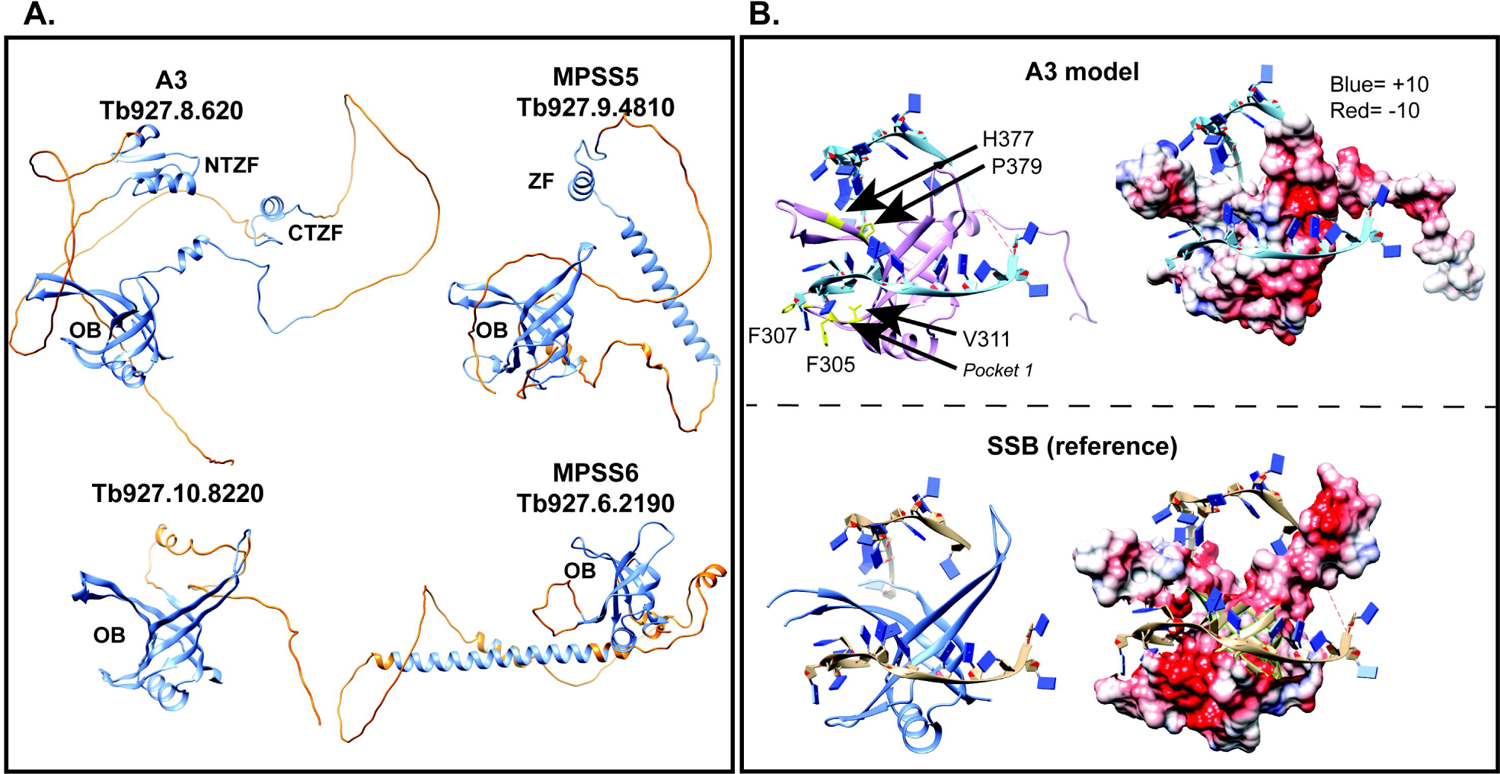
Structural similarities between A3 and other OB-fold proteins. **A**: AlphaFold predicted structures of A3, the predicted product from Tb927.10.8220, MPSS5, and MPSS6 showing the higher-(blue) and lower-(orange) -confidence (pLDDT<70) structures. OB-folds and the ZFs are noted. **B**: A3 model and SSB reference ribbon diagrams showing the L_12_ L_45_ groove (left) and Coulombic charge densities (units in kcal/mol*e) (right). Upper: Model of the A3 OB-fold interacting with ssDNA from the 3ULP crystal structure (PDBID: 4DNI). Lower: Chain C of the ssDNA bound SSB tetramer from *P. falciparum* (PDBID: 3ULP) used to make the model of A3. Locations of key residues in A3 are shown in yellow and the approximate location of a hydrophobic pocket identified by Park and Hol (2012) is indicated.

We developed a structural model to explore how A3 might bind RNA as many ZF and OB-fold containing proteins bind RNA or DNA and A3 has been suggested to bind RNA in vitro (Brecht et al. 2005; Voigt et al. 2018). A sequence and structural homology search using full-length A3 and HHpred (Zimmermann et al. 2018) identified the crystal structure of chain C of the tetrameric single-strand binding (SSB) apicoplast protein from *Plasmodium falciparum* as having the greatest structural similarity. Modeling of the A3 OB-fold (PDBID: 4DNI,) onto that of chain C in complex with ssDNA (PDBID: 3ULP) (Antony et al. 2012) which binds ss DNA via the groove between L_12_ and L_45_ shows how the A3 OB-fold might similarly bind RNA (Fig. 6B). The A3 groove appears narrower than that of SSB, but this may be due to conformational differences between RNA vs DNA binding. The calculated Coulombic charges of the L_12_ L_45_ groove of A3 is consistent with electrostatic interactions that could contribute to RNA binding and aromatic residues F305 and F307 in L_12_ are near modeled substrate implies potential base stacking interactions with RNA (Fig. 6B).

Secondary structures formed by RNA are more diverse than those formed by DNA, thus resulting in a diversity of mechanisms used by RNA-binding proteins to bind RNA (Corley et al. 2020), and CCs must accommodate a diverse set of substrates, editing sites, and mRNA structures. SSB was the closest structural homolog found but because it is a DNA binding protein we also searched the Protein Data Bank (PDB) for RNA-binding proteins with OB-folds complexed with RNA and found LIN28A. This protein interacts with a TUTase and regulates microRNAs (miRNAs) in higher eukaryotes (Wang et al. 2017). LIN28A contains an N-terminal cold-shock domain (CSD), which is a type of OB-fold, and two more C-terminal CCHC “zinc-knuckle” domains connected by flexible linkers which may provide flexibility for binding a variety of miRNA substrates (Nam et al. 2011; Wang et al. 2017). The IDRs in A3 likely provide a similar flexibility to accommodate the very numerous mRNA/gRNA substrates and edited sequences (Park and Hol 2012; Voigt et al. 2018), an observation that the AlphaFold model, which shows large unstructured regions, supports. These disordered regions could allow the three structured domains of A3 to coordinate during editing by providing flexibility to A3, and perhaps to the CCs by adopting dynamic conformations upon binding to other protein and/or RNA partners. A summary of findings showing relationships between CC structure and RNA editing is shown in Fig. 7.

**Figure 7:**
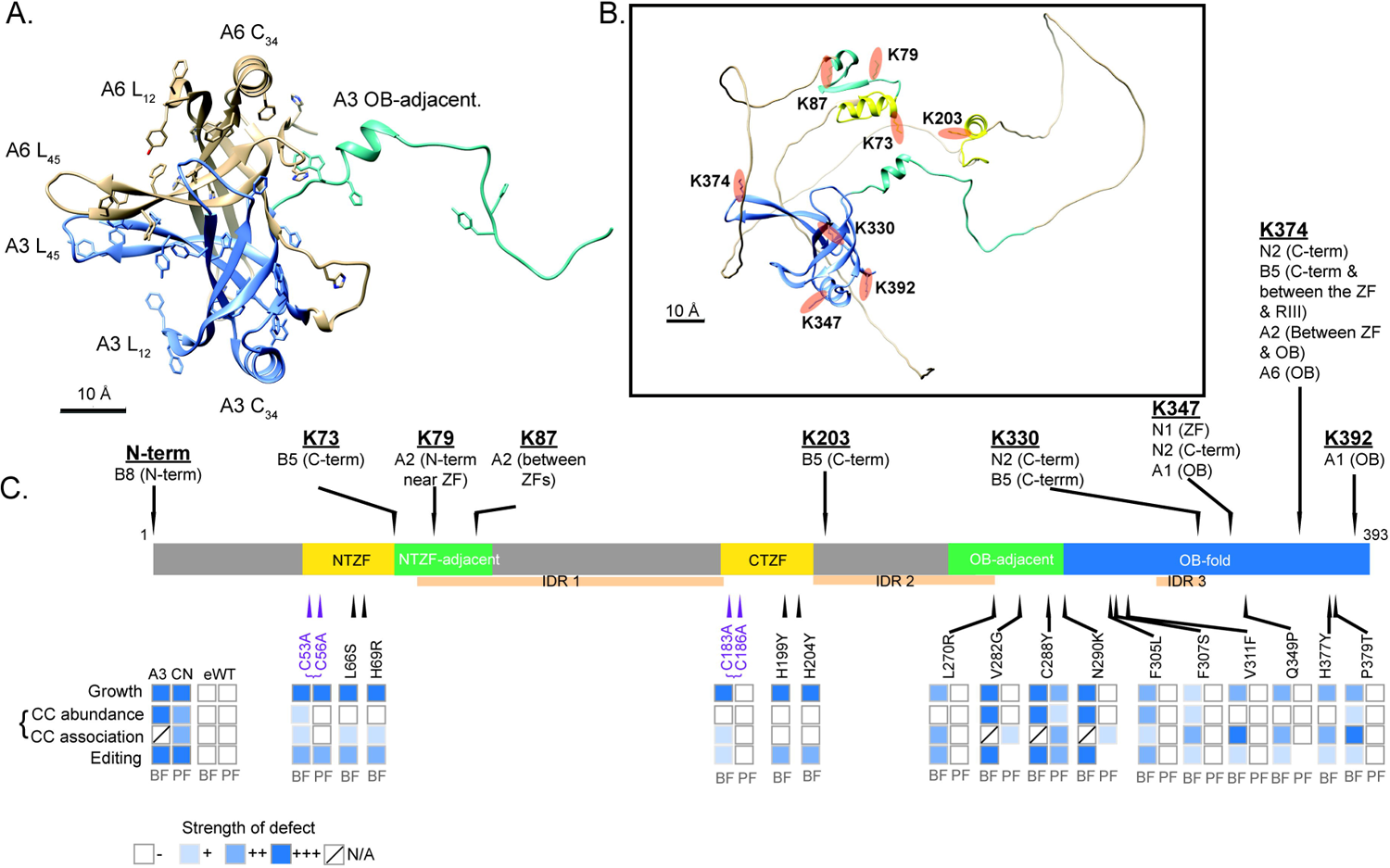
Summary of A3 protein characteristics, interactions and effects of mutations. **A**: Location of OB-fold domains and residues with planar side chains of the A3 OB-fold (blue) in complex with A6 (tan). **B**: Location of cross-linked Lys residues (orange) in the predicted AlphaFold model of A3 (McDermott et al 2016) showing the OB-fold (blue), ZFs (yellow) and ZF and OB-fold adjacent regions (teal). **C**: Summary diagram showing cross-links of A3 Lys residues with other CC proteins, domain locations, and effects of the mutations on cell line growth, CCs, and editing. Heat map indicates increasing strength of observed defects in CC abundance, association, and editing of A6, RPS12, and MURF2, as increasing intensity of blue shades (see legend).

## DISCUSSION

We show here that multiple functional domains of A3 are essential for structural integrity and accurate functioning of the CCs that perform U insertion and deletion editing. Multiple single amino acid (saa)-LOF mutations were identified in the two ZFs, C-terminal OB-fold, adjacent region domains and in a nearby intrinsically disordered region. Saa-LOF mutations that disrupt CC integrity indicate that A3 is critical to CC structure (Figs 2-5) and show that its interaction with A6 is especially important for this (Fig 7A). These results complement our previous results of A3 cross-linking to seven other CC proteins (summarized in Figs 7 and S3) (McDermott et al. 2016). The characteristics of the OB-fold and the results of mutations which alter but do not eliminate editing suggest functional interactions between A3 and substrate RNA. These may normally be associated with the accuracy and/or efficiency of editing, perhaps by affecting one or more of its many steps. The greater effects of A3 mutations on CC structure and editing in BFs than PFs mirror other studies which show that CCs are inherently different between life-cycle stages despite containing the same inventory of proteins (Carnes et al. 2011; McDermott et al. 2015b; Carnes et al. 2022). Our identification of structural homologs of A3 (Fig. 6 and Tables S3 and S4) may help illuminate the roles of these proteins that have functions associated with RNA.

### Multiple domains of A3 are critical for the structure and functions of the three different CCs

Each domain of A3 contributes to its overall function as is evident from the diverse effects of the mutations which range from the alteration but not loss of editing to the complete loss of CCs, their proteins and editing (summarized in Fig. 7). The C2H2 ZFs contribute to CC structural organization via their interactions with other CC proteins as shown by the effects of their mutation on CC integrity, i.e. less than ∼1 MDa CC material and reduced amounts of A1 protein (Fig 2), and the cross-linking of the ZFs with A2 and B5 (Schnaufer et al. 2010; McDermott et al. 2016). The greater effect of mutations of the N-terminal than the C-terminal ZF on CC structure suggests that this ZF and its adjacent fold participate in the protein interactions perhaps including proteins other than those found by cross-linking. That mutations of the ZFs alter but do not eliminate editing (Fig. 4) implies that they may also interact with RNA, perhaps transiently. The greatest impacts of ZF mutations that result from substitutions of the four C residues that coordinate the Zn^2+^ in both ZFs (Fig. 4) indicates that the A3 ZFs contribute to the network of functional protein and perhaps RNA interactions within CCs.

The A3 OB-fold contributes to CC structure and function via a variety of interactions with other CC proteins. The mutation analyses complement our previous cross-linking results which showed that the A3 OB-fold interacts with N1 and N2 in CC1 and CC2, respectively, and with A1, A6 and B5 that are in all CCs (Figs. 7 and S3) (McDermott, 2016). They extend understanding of the OB-fold interactions with A1 and A6 as V311F and P379T mutations resulted in loss of A1, and likely T2 and L2, other parts of the heterotrimeric insertion subcomplex (Schnaufer et al. 2003), and reduced CC abundance. V311 is part of a proposed hydrophobic pocket on the surface of A3 (Park and Hol 2012), and thus substitution to a bulky Phe may have disrupted this pocket suggesting that it participates in protein-protein interactions (Fig. 3). Most substitutions in or near β1 strand of the OB-fold (residues 282-305), part of which maps to the structural interface with A6 (Fig. 7)(Park and Hol 2012; Park et al. 2012), result in the absence of all CCs and their proteins (Figs. 2,3) indicating that the interaction between the A3 OB-fold and A6 is critical for CC structure. Thus, our results and those of others (Schnaufer et al. 2010; Park and Hol 2012; Park et al. 2012; McDermott et al. 2016) indicate that the OB-fold contributes to a network of protein-protein interactions within the CCs.

A3 contains three predicted IDR regions (Park and Hol 2012), but only the L270R mutation which is near an IDR was obtained in our random mutagenesis screen (Figs. 1 and 7). This may be because the mutagenesis was not extensive or due to the practical limitation on the number of cell lines characterized. The substantial impact of the L270R mutation on editing without an overall impact on CC structure suggests that it may have affected protein or RNA interactions and/or conformations within the CCs which are critical for editing. This implies that the other IDRs in A3 also have important functions in editing, perhaps by interacting with proteins or with RNA as suggested by the characteristics of these protein sequences which have been shown in other proteins to facilitate interactions with RNA (Zaharias et al. 2021; Zeke et al. 2022; Luo et al. 2023). The IDRs may provide the CCs with flexibility to accommodate many diverse substrate sequences and conformational changes during editing, like what has been shown for mammalian LIN28A (Nam et al. 2011; Wang et al. 2017).

### RNA interactions and effects on editing

Our structural modeling suggests that the A3 OB-fold can interact with RNA, i.e. substrate RNA (Fig. 6). The characteristics of the A3 OB-fold and its structural similarities to ssDNA and RNA-binding OB-fold proteins, e.g. *P. falciparum* SSB and mammalian LIN28A proteins (Theobald et al. 2003; Horvath 2011; Wang et al. 2017) suggests that these interactions could occur via a combination of base stacking interactions with planar residues and electrostatic interactions with a cluster of basic charges (Figs. 6 and 7). That several OB-fold mutations caused lesser effects on growth and CC structure and resulted in anomalous editing suggests that some of these mutations might have affected interactions between A3 and RNA. Given the interaction between A3 and A6, two or more OB-folds may cooperatively bind RNA, possibly via a mechanism analogous to the SSB tetramer binding of ssDNA (Antony et al. 2012). The data do not indicate which strand of RNA interacts with each OB-fold, but an attractive possibility is that one binds mRNA while the other binds gRNA. Alternatively, the OB-folds might work together to resolve secondary structures in the mRNA as has been previously suggested (Voigt et al. 2018). The editing substrate may also interact with other domains of A3 including its ZFs and IDRs both of which have been implicated in RNA binding in other systems (Tompa and Csermely 2004; Dyson 2012; Corley et al. 2020; Zaharias et al. 2021; Luo et al. 2023). Overall, the data suggest that during editing the substrate RNA interacts with A3 via base stacking and charged interactions within the OB-fold, and by interaction with certain residues within the IDRs and with the ZFs. Notably, substrate RNA is bound by the substrate complex (SC) during editing indicating that domains of CC proteins perhaps including those in A3 likely interact with SC proteins during editing.

### Developmental differences in editing

Differences in phenotypes resulting from the same substitutions in BF vs. PF indicate that A3 functions differently between the life cycle stages (Figs 3-6). There are several non-mutually exclusive possibilities that may explain the different BF vs. PF phenotypes: 1) A3 may undergo PTMs that are life-cycle stage specific, 2) A3 may interact with accessory factors that are present or function in editing in specific life-cycle stages, and 3) there may be conformational differences in BF vs. PF CCs, that result in different A3 interactions with other CC proteins in BF vs. PF. The results reported here mirror our previous results that show that CCs appear physically and functionally different between life-cycle stages despite having the same set of proteins (Carnes et al. 2011; Guo et al. 2012; McDermott et al. 2015a; McDermott et al. 2015b; McDermott and Stuart 2017; Carnes et al. 2022). The stabilities of CCs and CC proteins are more sensitive in BF relative to PF to most mutations or loss of CC proteins and most mutations of B5-B8 have greater effects on CC stability in BFs than PFs (Fig. 2) (Carnes et al. 2011; Guo et al. 2012; McDermott et al. 2015a; McDermott et al. 2015b; McDermott and Stuart 2017; Carnes et al. 2022). This implies that domain interactions may have lower affinities in BFs than PFs. Nevertheless, some mutations of CC proteins have the reciprocal life-cycle stage specific effects, i.e. detrimental in PF but not BF. These include the H233A mutation in the RAM motif of B5 that resulted in disruption of CCs and editing, and multiple mutations identified by deep mutational scanning of the RNase III domain of B4 (McDermott and Stuart, unpublished). The mechanisms underlying the differential editing are unknown (Schnaufer et al. 2002; McDermott et al. 2015b), but they likely involve interactions within CCs and between CCs, SCs and accessory factors that impact substrate specificities and kinetic characteristics of the multiple editing steps.

Overall, A3 appears to function in the structural organization and intramolecular positioning of proteins and substrate RNA within the three different CCs during editing. It may interact differently in CC1, CC2 and CC3 and affect ES binding and cleavage by their different endonucleases which initiates the editing of each ES and also differentially affect the post-cleavage steps in editing. The impacts of single amino acid substitutions throughout A3 on CC integrity and editing show that the region of A3/A6 interaction (Fig. 7A) (Wu et al. 2011; Park and Hol 2012; Park et al. 2012) is critical to CC stability and other regions are important for specific steps in the editing process. Structural modeling suggests how A3 might interact with RNA but further detailed analyses will be necessary to elucidate the specific effects of mutations in A3 on each of the three distinct CCs and on other components of the editing machinery, e.g. SCs. The structural homology between A3 and other proteins involved with mt RNA processing e.g. MPsome proteins (Fig 6, Tables S3 and S4) suggests that they have some similar functional characteristics which may aid elucidating their specific functions. Variants of the ZF, IDR and OB-fold domains of A3 function in a wide range of critical processes that entail protein and nucleic acid interactions which span DNA replication, repair and telomere maintenance as well as transcription, tRNA and microRNA processing, translation and others (Allison et al. 1998; Brevet et al. 2003; Theobald et al. 2003; Horvath 2011; Nam et al. 2011; Kapps et al. 2016; Wang et al. 2017; Amir et al. 2018; Gao et al. 2018). Results from this study may contribute the understanding of the roles of these domains in RNA editing and other processes.

## MATERIALS AND METHODS

### Preparation of pHD1344tub(PAC)-Nterm3V5-A3 and library construction

The A3 coding sequence minus the 20-codon predicted MTS (MitoFates (Fukasawa et al. 2015)) was amplified from the pHD1344-A3-myc plasmid (Guo et al. 2010) using primers that added AttB sites (Table S2). The resulting PCR product was first transferred into pDONR-221 via a BP clonase II reaction and then via an LR clonase II reaction into pHD1344tub(PAC)-Nterm3V5, which provides the MTS (Carnes et al. 2018). BF and PF A3 CN cells were transfected with pHD1344tub(PAC)-Nterm3V5-A3 and grown without tet to confirm expression and functional support of growth (Fig. 1). A mutagenized A3 library was produced and screened as previously described (Gray et al. 2007; McDermott et al. 2015a). Briefly, the A3 sequence specifying amino acids 21-393 was amplified from pHD1344tub(PAC)-Nterm3V5-A3 by error prone PCR using the GeneMorphII Random Mutagenesis kit (Agilent, Cat # 200523) and primers that add AttB sites (Table S2) from 1.4 µg of plasmid which had been determined to provide an optimal mutation rate. The mutated library was cloned into the pENTR-Express donor vector via a Gateway BP clonase II reaction (ThermoFisher), transformed into *E. coli* and plated on LB-agar containing 1mM IPTG and 40µg/mL kanamycin to eliminate frameshift and truncation mutants. The A3 stop codon that would interfere with the Kanamycin screening was eliminated during PCR amplification prior to library preparation.

The equivalent of ∼ 10,000 Kanamycin resistant clones were transferred into pHD1433tub(PAC)-Nterm3V5 via a Gateway LR-Clonase II reaction resulting in the pHD1344tub(PAC)-Nterm3V5-A3 mutagenesis library. The Gateway cloning of the mutated A3 sequence lacking a stop codon into pHD1344tub(PAC)-Nterm3V5-A3 resulted in an additional nine amino acids at the C-terminus of A3 that are absent in exclusively expressed WT A3 used in the A3 CN eWT cell line. The nine residues are present in the eWT allele used in the A3 CN+MGA eWT cell line. To assess the nature of the mutated A3 cloned library, 48 library clones were sequenced with primers 9571 and 11512 from which we estimate that 48.9% of the clones in the library have one or two amino acid changes, 25.5% have three or more changes and remaining 25.6% have no changes.

### Library screening

The mutated library was screened as previously described (McDermott et al. 2015a). The pHD1344tub(PAC)-Nterm3V5-A3 library was linearized with Not1, transfected into 1.5×10^8^ BF A3 CN cells (Guo et al. 2008), selected with 0.1µg/mL puromycin, and the cells were maintained in HMI-9 with penicillin-streptomycin, 10% FBS, and 5 ng/mL tetracycline, which is the minimum amount that allows normal cell growth. Cells were plated in 24 well plates at a density of ∼ one transfected cell per well which resulted in 634 puromycin-resistant cell lines. These potential cell clones were consolidated into seven 96 well plates, replica plated at a 1:100 dilution into media plus or minus tetracycline. The cells were grown for three days then passaged into another set of 96-well plates with plus or minus tet media for another three days. Following a total of six days of growth + or – tet, 20 μL of Alamar Blue cell viability reagent (Thermo Fisher) was added to each well and the plates were incubated for 4 h. Fluorescence was measured using a SpectraMax M2 microplate reader (Molecular Devices) with an excitation wavelength of 544nm and an emission wavelength of 590nm to identify wells with viable (pink) and non-viable cells (blue). Comparison of the tet plus and minus replica plates was used to identify 117 cell lines with a strong growth defect (blue in minus tetracycline) and 19 with a lesser growth defect (purple in minus tetracycline). These clones were consolidated into new 96 well plates and 10µL of confluent cells were lysed for gDNA extraction as previously described (McDermott et al. 2015a).

### PCR and sequencing of mutants

59 full growth defect clones were sequenced using primers 9571 and 5356 or 10150 (Tables S1 and S2). Forward and reverse sequence pairs were assembled using Geneious and aligned to the wild-type *T. brucei* 427 Lister A3 sequence to allow for identification of mutations and resulting amino acid substitutions (Table S1).

### Reconstruction of mutations and generation of exclusive expression cell lines

Mutations of interest, either single substitutions that disrupted function, substitutions found in more than one cell line, or substitutions at residues mentioned in the literature, were reconstructed in new cell lines as single amino acid substitutions using site-directed mutagenesis. The pHD1344tub(PAC)-Nterm3V5-A3 plasmid was mutagenized using the QuickChange Lightning site-directed mutagenesis kit (Agilent, product #210518) and primers listed in Table S2. All mutations were confirmed by Sanger sequencing using primers 9571 and 5356 or 10150 (Table S2). Substitutions of the C residues in both ZF domains were made using site-directed mutagenesis and the primers listed in Table S2. The N-terminal ZF (NTZF), contains both C53A and C56A, substitutions while the more C-terminal ZF (CTZF) contains C183A and C186A, and the NTZF&CTZF plasmid contains all four C➔A mutations as previously described (Guo et al. 2008; Guo et al. 2010). Our new V5-tagged ZF constructs were used for our study as no protein could be detected from a previous construct, NTZF-myc, as previously described (Fig. S4) (Guo et al. 2010). Plasmids were linearized using Not1 and transfected into the BF A3 CN cell line and selected with 0.1µg/mL puromycin as previously described (Guo et al. 2008; Merritt and Stuart 2013). The resulting puromycin-resistant cell lines were screened using PCR with primers 5355 and 10150 verify integration of the plasmid into the tubulin locus, and Western blot to check for expression of V5-tagged A3. Plasmids encoding mutations resulting in BF growth defects were also transfected into PF A3 CN cells as previously described, except for H377Y and mutations in the ZFs (Guo et al. 2008; Guo et al. 2010; McDermott et al. 2015b). PF transfectants were genotyped with PCR as above using primers 5355 and 10150, and cell lysates and expression of the V5-tagged A3 was confirmed by western blot (data not shown).

### Generation of A3 mutants in the BF A3 CN+MGA cell line background

The BF A3 CN+ MGA cell line was made by transfecting the A3 CN cells (Guo et al. 2008) with pEnT6+ATPaseGammaWT+3UTR that carries the L262P mutation in gamma ATPase, and selected with blasticidin (Dean et al. 2013; Carnes et al. 2017). The resulting cell line, A3 CN+ MGA, was genotyped by PCR and sequenced using primers 9530 and 9531 (Table S2) to confirm the presence of the MGA mutation, and rescue of the A3 CN growth defect in the absence of tetracycline was confirmed. The same pHD1344tub(PAC)-Nterm3V5-A3 plasmids were transfected into the A3 CN+MGA cell line as previously described (Guo et al. 2010; Merritt and Stuart 2013).

### Cell culture and transfections

BF cells were grown in HMI-9 with 10% FBS at 37 °C, 5% CO2. PF cells were grown in SDM-79 with 10% FBS at 27 °C. The PF A3 CN cell line used contained an untagged tet-regulated A3 construct expressed from the rDNA locus and was produced as previously described (McDermott et al. 2015b). Unless otherwise stated, the concentrations of drugs used for selection and tet-regulated expression of transgenes in this study were as follows: for BFs, 2.5 μg/ml G418, 5 μg/ml hygromycin, 2.5 μg/ml phleomycin, 0.5 μg/ml Tet, and 0.1 μg/ml puromycin; for PFs, 15 μg/ml G418, 25 μg/ml hygromycin, 2.5 μg/ml phleomycin, 0.5 μg/ml Tet, 1 μg/ml puromycin, and 10 μg/ml blasticidin. For BF transfections, cells were washed in PBS plus 6mM glucose, resuspended in 0.2M Sodium phosphate, 2M KCl, 0.1M CaCl2, and 2M HEPES, and transfected using the Amaxa nucleofector (Lonza) program X-001 as previously described (Merritt and Stuart 2013). For PF transfections, cells were washed and resuspended in 0.5mL cytomix buffer (25mM HEPES pH 7.6, 120mM KCl, 0.15mM CaCl2, 10mM K2HPO4/KH2PO4 pH 7.6, 2mM EDTA, 6 mM glucose, 5mM MgCl2. Washed cells were then transfected using a BTX nucleofector at 1600V, 25Ω, 16µF and plated as previously described (Merritt and Stuart 2013).

### Growth rate analyses

BF cells were seeded at an initial density of 2×10^5^ cells/mL and PF cells at 2×10^6^ cells/mL unless noted otherwise. Cells were counted using a Z1 Coulter particle counter each day. BF were reseeded at 2 x 10^5^ cells/mL in 10 mL every day, while PF were reseeded at 2 x 10^6^ cells/mL in 10 mL every two days. The ratio of cumulative cell numbers in -tet to + tet cultures for each cell line was calculated (Carnes et al. 2022). Log2 of the ratio is displayed on a blue-orange heat map created using Microsoft Excel.

### Glycerol gradient fractionation

Glycerol gradients were run as previously described (McDermott et al. 2015a). Briefly, 3×10^9^ BFs were grown in the absence of tet for 72 hours, collected by centrifugation and washed with PBS plus 6mM glucose. Cell pellets were flash frozen in liquid nitrogen and stored at −80°C until use. Pellets were lysed in 500-1000µL of lysis buffer containing 10mM Tris pH 7.2, 10mM MgCl_2_, and 100mM KCl supplemented with cOmplete mini protease inhibitor tablets (Roche), pepstatin A, leupeptin, Pefabloc (Roche), and 1 mM DTT. Triton-X100 was then added to a final concentration of 1%. Cells were incubated on ice for 10 min followed by centrifugation at 13,000 rpm in a microcentrifuge to clear. 50 to 100µL of the cleared lysate was taken for a western blot and the remainder was loaded onto 11 mL 10-30% glycerol gradients and centrifuged in a Beckman Optima XPN-80 ultracentrifuge for 9 hours at 38,000 rpm using an SW40 rotor. 500µL fractions were collected from the top and 100µL samples were taken for analysis by both native and denaturing western blots. Remaining fractions were flash frozen in liquid nitrogen and stored at −80°C. Western blots from gradients were developed using either ECL (Pierce) or SuperSignal West PICO Plus ECL (Pierce).

### SDS-PAGE, BN-PAGE, and Western blotting

Western blots were conducted as previously described (Carnes et al. 2022). Briefly, cell pellets containing 5×10^7^-2×10^8^ cells were resuspended on ice in IPP150 buffer (10mM Tris-HCL pH 8.0, 150mM NaCl, and 0.1% NP-40) with cOmplete protease-inhibitor cocktail mini tablets (Roche). Triton-X100 was added to a final concentration of 1% and the samples were incubated on ice for 10-20 mins to lyse the cells. Lysates were cleared by centrifugation at 13,000 rpm in a microcentrifuge for 10 mins, diluted 1:1 in Tricine sample buffer (Bio Rad) with β-mercaptoethanol and up to ∼6×10^6^ cell equivalents were loaded onto each lane of 10% TGX-acrylamide Criterion gels (Bio Rad). Gels were run at 100V for ∼ 2 h or until the dye front ran off the gel. Protein was transferred to PVDF membrane (Millipore) using a Tris/glycine transfer buffer with 20% methanol. Membranes were blocked in PBS-Tween with 5% milk for 15 min before the addition of primary antibody. Antibodies and dilutions are listed in Table S4. Western blots were exposed to ECL (Pierce) and visualized using X-ray film (McKesson). For BN-PAGE, cleared lysates were prepared the same as for the denaturing western blots. 5×10^6^ cell equivalents were loaded into wells on a 3-12% Bis-Tris gradient gel with unstained NativePage protein standards in the first well (ThermoFisher) and run according to manufacturer’s instructions. Protein was then transferred to a PVDF membrane (Millipore) overnight with NuPAGE transfer buffer ThermoFisher) without methanol at 22-24V at 4°C. Following transfer, gels were stained with Ponceau SP to visualize the protein standards, fixed in 10% acetic acid, and probed with antibodies (Table S6).

### RNA extraction, RT-PCR and qPCR

All procedures for RNA extraction, quantification, cDNA synthesis, and qPCR were designed to follow the MIQE guidelines. Briefly, 1×10^8^ cells were harvested by centrifugation, washed with PBS and 6mM glucose, resuspended in Trizol, and stored at −80°C until use. Total RNA was extracted from the Trizol samples, quality was checked using a BioAnalyzer (Agilent) and quantified using the NanoDrop One spectrophotometer (Thermo Scientific). RNAs were treated with DNase I and cDNA was generated using random hexamer primers with TaqMan reverse transcription reagents and MultiScribe reverse transcriptase (ThermoFisher). Prior to qPCR, cDNAs were pre-amplified using Taqman PreAmp master mix (ThermoFisher), treated with Exonuclease I to remove excess primers, and diluted as previously described (McDermott et al. 2015a). High-throughput BioMark qPCR was performed as previously described using SsoFast EvaGreen with low ROX (Bio-Rad) and the Fluidigm BioMark HD system (McDermott et al. 2015b). Data was analyzed using the Fluidigm real-time PCR analysis software and Microsoft Excel. In most cases, heat maps were generated using the average of two or three biological replicates with two technical replicates each. However, there is only one ND7 3’ ed and ND8 ed replicate for V311F, H377Y, and P379T. Primers used for this assay are listed in Table S2. For RT-PCR (Fig. 4), cDNA was synthesized from 1µg of RNA using Multiscribe reverse transcriptase (Invitrogen) and a gene specific reverse primer (4394 (MURF2), 5380 (A6), 3620 (RPS12), or 3707 (ND4)) (Table S2). The entire amount of cDNA was used in a PCR reaction to amplify MURF2, RPS12, A6, or ND4 transcripts using primer pairs 6204/4934 (MURF2), 3704/3580 (A6), 3619/3620 (RPS12), and 3706/3707 (ND4) (Table S2) (Carnes et al. 2022). PCR products were resolved on a 3% agarose gel containing ethidium bromide and visualized.

### Cloning and Sanger sequencing of RT-PCR products

Remaining RT-PCR products from Figure 4 were resolved on an 1.5% agarose gel and all RPS12 products spanning pre-edited through edited sizes (∼200-350 bp) were excised for each sample (BF L270R, BF eWT, and BF CN), purified, and cloned into pGEM-T-Easy (Promega) according to manufacturer’s instructions. Ligations were transformed into 5α *E. coli* (NEB) and plated on Amp/IPTG/X-gal plates for blue/white screening. White colonies were selected for Sanger sequenced using T7 and M13R primers and assembled using Geneious. 11 clones for eWT, 24 clones for L270R, 13 clones for F307S, and 16 clones for P379T were analyzed. 4 clones from the CN were also obtained. Editing events were determined and sequences aligned by hand. Editing sites were defined as each position between two non-U nucleotides, beginning with the most 3’ deletion site in RPS12, as this region was not spanned by primer 3620 used for RT-PCR. Editing at each site was then tabulated for each set of samples and recorded as follows:: I-Ed.= expected insertion sites that are edited (the number of Us matches the fully edited (F.E.) RPS12 sequence), I-Part. Ed.= partially-edited insertion sites (do not match F.E. or Pre-edited (P.E.) RPS12), D-Ed.= edited deletion sites, D-Part. Ed.= partially-edited deletion sites, N-Part. Ed.= partial editing at a site that is non edited in F.E. RPS12, not edited, and N/A= a clone contained an A, C, or G mismatch so it could not be analyzed at one or more ESs. Graphs, which show the type of editing at each site as a percent of total transcripts, were generated using Microsoft Excel. Graphs include all potential ESs (positions between non-U nucleotides). The type of editing (insertion, deletion, or no editing) expected at each site based on the fully-edited RPS12 sequence is indicated beneath each bar by a yellow (insertion), blue (deletion), or black (no editing) dot. Aligned sequences from each set of samples are shown in Figure S2.

### Protein structure modeling

Full-length A3 was searched using the HHpred structural prediction tool (https://toolkit.tuebingen.mpg.de/tools/hhpred). Structures of A3 bound to nucleic acid were modeled using HHpred and Modeller, and all protein structures and models were visualized using Chimera (Fiser and Sali 2003; Pettersen et al. 2004; Zimmermann et al. 2018). A3 sequence was modeled onto the structure of *Plasmodium falciparum* SSB (single-strand binding protein) in complex with ssDNA, PDB ID: 3ULP (Antony et al. 2012). 3ULP was chosen for the nucleic-acid bound OB-fold model as HHpred listed it as the most similar OB-fold to the query sequence, with a probability of 97.8% and E=0.0011. Five models were calculated using the Modeller program and the highest confidence model (model 2.4, not shown), with GA341 and zDOPE scores of 1.0 and −0.08, respectively, was chosen for further analysis. Hydrophobicity and Coulombic charges were calculated by Chimera (Pettersen et al. 2004). The predicted structure of the *T. brucei* A3, MPSS5, MPSS6, and the protein product encoded by Tb927.10.8220 (Uniprot IDs D6XMN0, Q38F66, Q584T7, and Q38AE2, respectively) shown in Figures 1 and 6 and the cross-linking partners of A3 shown in figure S3 (N1=Q4GZ50, N2=Q38B60, B8=Q57X13, A1=Q586L9, A2=Q38AE3, B5=Q387F6) were obtained from the AlphaFold Protein Structure Database version 2022-11-01, created with the AlphaFold Monomer v2.0 pipeline (https://alphafold.ebi.ac.uk/) (Jumper et al. 2021; Varadi et al. 2022). A Dali structural homology search was conducted using A3 (D6XMN0) (http://ekhidna2.biocenter.helsinki.fi/dali/) (Holm 2022). Only proteins that may localize to the mitochondrion are listed in table S3. Potential cognate residues in proteins with structural similarity to A3 (Table S4) were visually identified using the AlphaFold structures of each protein in comparison to A3.

### Multiple sequence alignments

Sequence alignments in Figure 1 were constructed using the MUSCLE alignment tool in Geneious. The gene IDs for *T. brucei*, *T. cruzi*, and *T. congolense* A3 used for the alignment are Tb427.08.620, TcCLB.509611.110, and TcIL3000_8_100.1, respectively.

## ACKNOWLEDGMENTS

This work was supported by the National Institutes of Health grant R01AI014102 to K.S. We thank Xuemin Guo for providing the A3 CN cell lines.

---

**Figure S1: Parasite growth and CC integrity in BF MGA cell lines A**: Native western blots of MGA cell lines after 48 and 96 hours since tetracycline withdrawal. Western blots were probed with the A2 mAb to visualize CCs. **B**: Growth data for BF A3 mutant cell lines carrying the MGA mutation. Blue indicates fewer cells in the –tet sample compared to + tet, whereas orange indicates more cells in the –tet sample.

**Figure S2: RPS12 sequence characteristics from different BF cell lines**. Sequences from each cell line were aligned with each other and fully edited and pre-edited RPS12. Inserted and deleted Us that match the F.E. sequence are indicated in red or by *, respectively. Inserted and deleted Us that do not match the F.E. sequence are also indicated. Junctions in each transcript are underlined. Highlighted regions contain a non-U nucleotide mismatch. Analysis of this data to produce the graphs in Figure 5 is described in detail in the Materials and Methods.

**Figure S3: Summary of A3 cross-links within CCs A**: Summary of the lysine residues in A3 that were involved in cross-links as described by McDermott et al (2016). Cross-linked lysine residues in A3 are indicated in orange. **B-H**: Locations of cross-links shown on structural diagrams (AlphaFold) of A3 and each cross-linked protein.

**Figure S4: Comparison between V5-NTZF and NTZF-myc cell lines.** Denaturing western blot using mAbs to detect four CC proteins (A1, A2, L1, and A3) in exclusively-expressed A3 cell lines with either V5 or myc tags. No A3 protein is observed in the NTZF-myc sample.

